# Top-down task goals induce the retrieval state

**DOI:** 10.1101/2024.03.04.583353

**Authors:** Devyn E. Smith, Nicole M. Long

**Affiliations:** Department of Psychology, University of Virginia 22904

**Keywords:** encoding, retrieval, brain state, EEG, MVPA

## Abstract

Engaging the retrieval state (Tulving, 1983) impacts processing and behavior (Long & Kuhl, 2019, 2021; Smith, Moore, & Long, 2022), but the extent to which top-down factors – explicit instructions and goals – vs. bottom-up factors – stimulus properties such as repetition and similarity – jointly or independently induce the retrieval state is unclear. Identifying the impact of bottom-up and top-down factors on retrieval state engagement is critical for understanding how control of task-relevant vs. task-irrelevant brain states influence cognition. We conducted between-subjects recognition memory tasks on male and female human participants in which we varied test phase goals. We recorded scalp electroencephalography and used an independently validated mnemonic state classifier (Long, 2023) to measure retrieval state engagement as a function of top-down task goals (recognize old vs. detect new items) and bottom-up stimulus repetition (hits vs. correct rejections). We find that whereas the retrieval state is engaged for hits regardless of top-down goals, the retrieval state is only engaged during correct rejections when the top-down goal is to recognize old items. Furthermore, retrieval state engagement is greater for low compared to high confidence hits when the task goal is to recognize old items. Together, these results suggest that top-down demands to recognize old items induce the retrieval state independent from bottom-up factors, potentially reflecting the recruitment of internal attention to enable access of a stored representation.

**Significance Statement:** Both top-down goals and automatic bottom-up influences may lead us into a retrieval brain state – a whole-brain pattern of activity that supports our ability to remember the past. Here we tested the extent to which top-down vs. bottom-up factors independently influence the retrieval state by manipulating participants’ goals and stimulus repetition during a memory test. We find that in response to the top-down goal to recognize old items, the retrieval state is engaged for both old and new probes, suggesting that top-down and bottom-up factors independently engage the retrieval state. Our interpretation is that top-down demands recruit internal attention in service of the attempt to access a stored representation.

## Introduction

Despite evidence that a tonically maintained retrieval state (or mode; Tulving, 1983) impacts behavior (Long & Kuhl, 2019), the factors that govern how the retrieval state is engaged are unclear. Although considered a goal-driven, intentional state (Rugg & Wilding, 2000), the retrieval state may also be induced automatically (Duncan, Sadanand, & Davachi, 2012; Smith et al., 2022). If asked if you have seen the film *Best in Show*, you may turn your mind’s eye inward and conclude that you have not. Alternatively, after hearing the title, you might automatically have images brought to mind of Parker Posey melting down in a pet shop. In the first example, top-down demands (try to retrieve *Best in Show*) may lead you to intentionally engage the retrieval state. In the second example, bottom-up signals – activation of a stored representation – may automatically pull you into the retrieval state. As these factors are typically considered in isolation, the relative contribution of bottom-up and top-down factors to retrieval state engagement remains an open question. Addressing this question is important as top-down vs. bottom-up driven states likely recruit distinct control mechanisms. The aim of this study was to identify the joint impact of bottom-up and top-down factors on retrieval state engagement.

The retrieval state may constitute an intentional state that is a precursor to successful retrieval (Lepage, Ghaffar, Nyberg, & Tulving, 2000; Herron & Wilding, 2004, 2006). An individual may explicitly engage a brain state – a collection of whole-brain activity and connectivity patterns sustained over time (Harris & Thiele, 2011) – when given the goal to retrieve and prior to accessing a stored representation. Using an explicit mnemonic state task in combination with multivariate decoding methods, we have shown that individuals can flexibly engage the retrieval state and that whole-brain spectral patterns dissociate the retrieval state from memory encoding (Smith et al., 2022). Retrieval state engagement influences how stimuli are processed (Long & Kuhl, 2021) and later remembered (Long & Kuhl, 2019). Thus, retrieval has downstream consequences, yet the question of how bottom-up factors govern retrieval state engagement remains open.

Bottom-up factors can induce retrieval independent from top-down demands (Duncan et al., 2012; Smith et al., 2022). Behavioral evidence from continuous recognition paradigms suggests that perceiving an item as old may induce a lingering retrieval state that influences subsequent decisions (Duncan et al., 2012). Judging a probe as “old” vs. “new” may serve to bias the hippocampus towards either pattern completion or pattern separation (O’Reilly & McClelland, 1994; Yassa & Stark, 2011). Similarly, temporal overlap between non-identical stimuli can induce retrieval (Smith et al., 2022). These results suggest a role for bottom-up factors in retrieval state engagement.

Insofar as the retrieval state reflects internally directed attention (Chun, Golomb, & Turk-Browne, 2011; Long, 2023), bottom-up vs. top-down factors should differentially modulate when and how this state is engaged. Top-down demands will recruit internal attention to enable representation access and should thus emerge earlier than bottom-up induced retrieval, which should occur in response to the access attempt. Similarly, difficult-to-access representations should increase top-down driven retrieval whereas easy-to-access representations should increase bottom-up driven retrieval, whereby internal attention is captured by accessed content (Cabeza et al., 2011). This framework suggests that top-down driven retrieval should emerge whenever there are explicit retrieval demands.

Our hypothesis is that top-down and bottom-up factors do not interact and specifically that top-down demands fully engage the retrieval state. Alternatively, both factors may be additive such that top-down demands coupled with stimulus repetition produce a stronger retrieval state than either factor alone. To test our hypothesis, we conducted independent recognition memory studies in which we varied top-down test phase goals while recording scalp electroencephalography (EEG). We used cross-study classification to measure test phase retrieval state engagement. We predicted that retrieval state evidence would increase for all test trials when task demands required participants to recognize old stimuli and stored representations were difficult to access.

## Materials and Methods

### Participants

Seventy six native English speakers from the University of Virginia community participated, with thirty eight participants enrolled in each experiment (E1: 28 female; age range = 18-32, mean age = 20.47 years; E2: 26 female; age range = 18-32, mean age = 20.5 years). All participants had normal or corrected-to-normal vision. Informed consent was obtained in accordance with University of Virginia Institutional Review Board for Social and Behavioral Research and participants were compensated for their participation. Our sample size was determined *a priori* based on behavioral pilot data (E1, N = 5; E2, N = 3) described in the pre-registration report of this study (https://osf.io/6u9px). A total of four participants (two each from E1 and E2) were excluded from the final dataset due to EEG event markers not being recorded. Thus data are reported for the remaining seventy two participants.

### Experimental Design

We conducted two recognition memory experiments (E1, E2) and manipulated test phase instructions between subjects. Participants’ goal was to successfully retrieve study items (E1) or to detect new items (E2). Stimuli consisted of 1602 words, drawn from the Toronto Noun Pool (Friendly, Franklin, Hoffman, & Rubin, 1982). From this set, 640 words were randomly selected for each participant. Participants completed four phases (Figure 1). Phase 3 was divided into two subsets; the practice subset preceded Phase 1 and the main subset preceded Phase 3.

**Figure 1.**
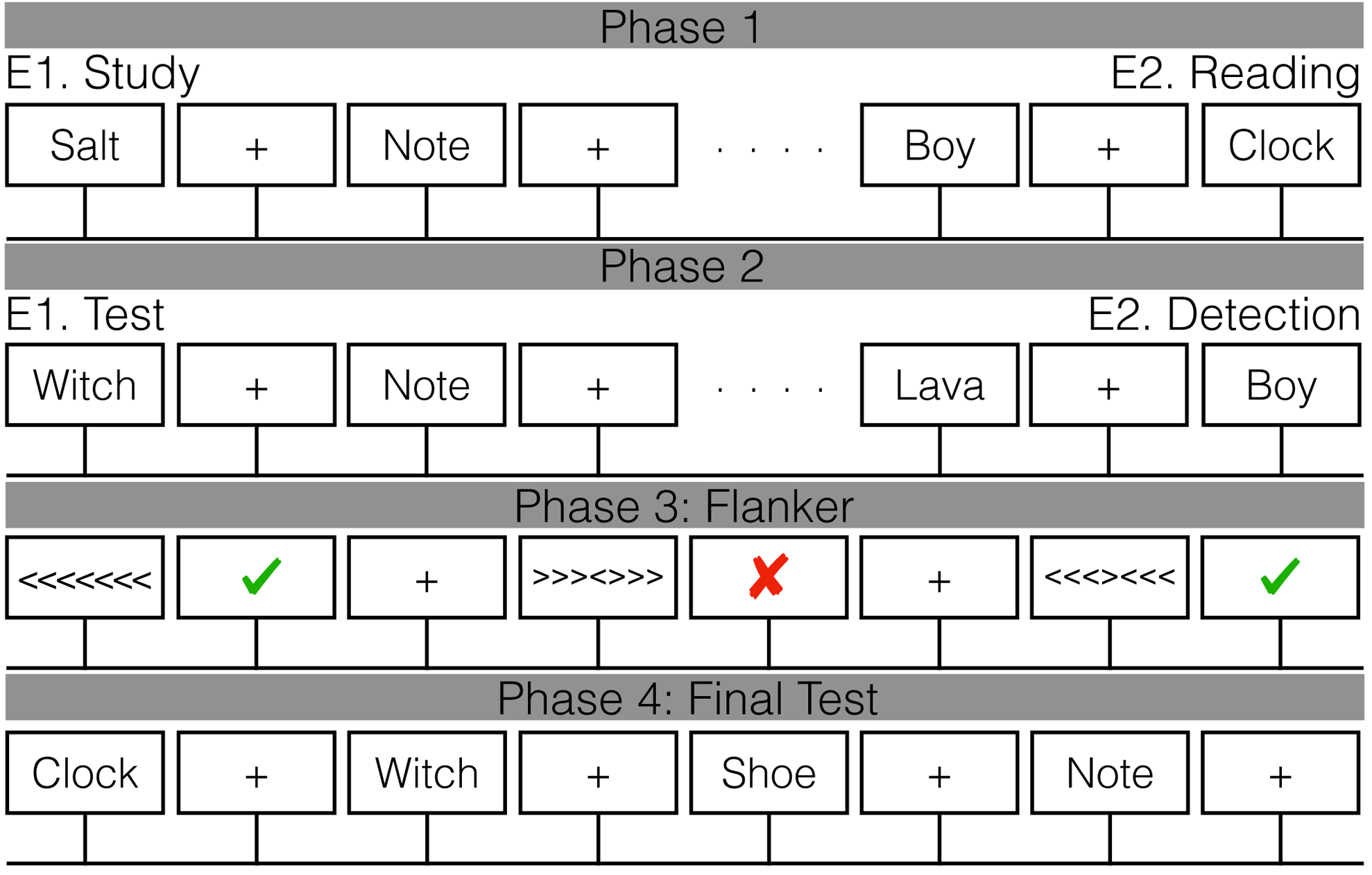
Task design. The Phase 3 flanker task was divided into a practice subset of three runs and a main subset of six runs. The practice subset was completed prior to Phase 1 to determine response duration during the main subset (see Methods). In E1 Phase 1, participants studied individual words in anticipation of a later memory test. In E2 Phase 1, participants read the words silently. In E1 Phase 2, participants completed a recognition test and made old or new judgements using a confidence rating scale from 1 to 4, with 1 being definitely new and 4 being definitely old. In E2 Phase 2, participants completed a detection phase in which the goal was to detect new words that were not presented in Phase 1. Participants made old or new judgements without the use of a confidence rating scale. All participants then completed Phase 3, a flanker task, in which they made speeded responses to a central target. Immediately after each response, a green check mark indicating a correct response or a red X indicating an incorrect response was presented. In Phase 4, participants completed a final recognition memory test in which all the words from Phases 1 and 2 were presented along with novel lures. Participants made old or new judgements using a confidence rating scale from 1 to 4, with 1 being definitely new and 4 being definitely old.

*Phase 1.* In each of 10 runs, participants viewed a list containing 16 words, yielding a total of 160 trials. On each trial, participants saw a single word presented for 2000 ms followed by a 1000 ms inter-stimulus interval (ISI). In E1, participants were instructed to study the presented word in anticipation for a later memory test and did not make any behavioral responses. In E2, participants were instructed to read the words silently and did not make any behavioral responses.

*Phase 2.* Participants completed a recognition memory test with different memory goals. On each trial, participants viewed a word which had either been presented during Phase 1 (target) or had not been presented (lure). In E1, participants’ task was to make an old or new judgement for each word using a confidence rating scale from 1 to 4, with 1 being definitely new and 4 being definitely old. In E2, the task was framed as a detection phase in which participants’ task was to detect new words that were not presented in Phase 1. Participants made an old or new judgment without the use of a confidence rating scale for each word by pressing one of two buttons (“d” or “k”). Response mappings were counterbalanced across participants. Phase 2 trials were self-paced and separated by a 1000 ms ISI. There were a total of 320 test trials with all 160 Phase 1 words presented along with 160 novel lures.

*Phase 3.* Prior to beginning Phase 1, participants completed three practice runs of a flanker task in which they made speeded responses to a central target in a string of congruent (e.g. >>>>>>>) or incongruent (e.g. <<<><<<) arrows. Feedback was presented immediately after each response as either a green check mark indicating a correct response or a red X indicating an incorrect response. Response duration, the interval in which a response was accepted, was initially set to 375 ms based on pilot data. To maintain difficulty and ensure an approximately balanced number of correct and incorrect responses during the main subset of Phase 3, response duration was individually adjusted based on participants’ accuracy following each practice run. If accuracy was below 50%, response duration increased by 25 ms, if accuracy was above 50%, response duration decreased by 25 ms. Thus, after completing the three practice runs, the final response duration could be a minimum of 300 ms and a maximum of 450 ms. After completing Phase 2, participants completed the main subset of Phase 3 which consisted of six runs of the flanker task. Throughout the main subset of Phase 3, the response duration was fixed to that obtained from the final practice run. As our analyses focus on Phase 2, we do not consider the flanker data further.

*Phase 4.* Participants completed a final recognition memory test in which all the words from Phase 1 and 2 were presented along with novel lures. Trials were self-paced and participants made old or new judgements for each word using a confidence rating scale from 1 to 4, with 1 being definitely new and 4 being definitely old. Trials were separated by a 1000 ms ISI. There were a total of 640 test trials with all 320 Phase 2 words presented along with 320 novel lures. As our analyses focus on Phase 2, we do not consider the final test data further.

### Independent Recognition Memory Experiment

To control for differing demands with regards to confidence judgments, we also include data from a third recognition memory experiment (“E3”) on which we have previously reported (Smith, Wheelock, & Long, 2024). All of the analyses and results described here are new. Briefly, in E3, participants studied twelve sixteen-item word lists in anticipation of a later memory test. During the test phase, participants made old/new recognition memory judgments to targets and lures.

### Mnemonic state task

An independent group of participants completed a mnemonic state task. Participants were biased via explicit instructions on a trial-by-trial basis to engage an encoding or retrieval state, while perceptual input and behavioral demands were held constant. In this mnemonic state task (for specific study parameters, please see refs. Smith et al., 2022; Hong, Smith, Moore, & Long, 2023), participants viewed two lists of object images. For the first list, each object was new. For the second list, each object was again new but was categorically related to an object from the first list. For example, if List 1 contained an image of a bench, List 2 would contain an image of a different bench. During List 1, participants were instructed to encode each new object. During List 2, however, each trial contained an instruction to either encode the current object (e.g., the new bench) or to retrieve the corresponding object from List 1 (the old bench). Each object was presented for 2000 ms. Participants completed either a two-alternative forced choice recognition test or a recency test on the object stimuli. We used the stimulus-locked List 2 data to train a multivariate pattern classifier (see below) to distinguish encoding and retrieval states.

### EEG Data Acquisition and Preprocessing

EEG recordings were collected using a BrainVision system and an ActiCap equipped with 64 Ag/AgCl active electrodes positioned according to the extended 10-20 system. All electrodes were digitized at a sampling rate of 1000 Hz and were referenced to electrode FCz. Offline, electrodes were later converted to an average reference. Impedances of all electrodes were kept below 50kΩ. Electrodes that demonstrated high impedance or poor contact with the scalp were excluded from the average reference. Bad electrodes were determined by voltage thresholding (see below).

Custom python codes were used to process the EEG data. We applied a high pass filter at 0.1 Hz, followed by a notch filter at 60 Hz and harmonics of 60 Hz to each participant’s raw EEG data. We then performed three preprocessing steps (Nolan, Whelan, & Reilly, 2010) to identify electrodes with severe artifacts. First, we calculated the mean correlation between each electrode and all other electrodes as electrodes should be moderately correlated with other electrodes due to volume conduction. We z-scored these means across electrodes and rejected electrodes with z-scores less than −3. Second, we calculated the variance for each electrode, as electrodes with very high or low variance across a session are likely dominated by noise or have poor contact with the scalp. We then z-scored variance across electrodes and rejected electrodes with a |z| > = 3. Finally, we expect many electrical signals to be autocorrelated, but signals generated by the brain versus noise are likely to have different forms of autocorrelation. Therefore, we calculated the Hurst exponent, a measure of long-range autocorrelation, for each electrode and rejected electrodes with a |z| > = 3. Electrodes marked as bad by this procedure were excluded from the average re-reference. We then calculated the average voltage across all remaining electrodes at each time sample and re-referenced the data by subtracting the average voltage from the filtered EEG data. We used wavelet-enhanced independent component analysis (Castellanos & Makarov, 2006) to remove artifacts from eyeblinks and saccades.

#### EEG Data Analysis

We applied the Morlet wavelet transform (wave number 6) to the entire EEG time series across electrodes, for each of 46 logarithmically spaced frequencies (2-100 Hz; Long & Kahana, 2015), across all experiments. After log-transforming the power, we downsampled the data by taking a moving average across 100 ms time intervals from −1000 to 3000 ms relative to the response and sliding the window every 25 ms, resulting in 157 time intervals (40 non-overlapping). Mean and standard deviation power were calculated across all trials and across time points for each frequency. Power values were then z-transformed by subtracting the mean and dividing by the standard deviation power. We followed the same procedure for the mnemonic state task, with 317 overlapping (80 non-overlapping) time windows from 4000 ms preceding to 4000 ms following stimulus onset (Smith et al., 2022).

#### Pattern Classification Analyses

Pattern classification analyses were performed using penalized (L2) logistic regression implemented via the sklearn module (0.24.2) in Python and custom Python code. For all classification analyses, classifier features were comprised of spectral power across 63 electrodes and 46 frequencies. Before pattern classification analyses were performed, an additional round of z-scoring was performed across features (electrodes and frequencies) to eliminate trial-level differences in spectral power (Kuhl & Chun, 2014; Long & Kuhl, 2018; Smith et al., 2022). Therefore, mean univariate activity was matched precisely across all conditions and trial types. We extract “classifier evidence,” a continuous value reflecting the logit-transformed probability that the classifier assigns the correct mnemonic label (encode, retrieve) for each trial. Classifier evidence is used as a trial-specific, continuous measure of memory state information, which is used to assess the degree of retrieval state evidence present during hit and correct rejection trials.

#### Cross study memory state classification

To measure retrieval state engagement in E1, E2 and E3, we conducted three stages of classification using the same methods as in our prior work (Long, 2023). First, we conducted within participant leave-one-run-out cross-validated classification (penalty parameter = 1) on all participants who completed the mnemonic state task (N = 100, see ref. Hong et al., 2023 for details). The classifier was trained to distinguish encoding vs. retrieval states based on spectral power averaged across the 2000 ms stimulus interval during List 2 trials. For each participant, we generated true and null classification accuracy values.

We permuted condition labels (encode, retrieve) for 1000 iterations to generate a null distribution for each participant. Any participant whose true classification accuracy fell above the 90th percentile of their respective null distribution was selected for further analysis (N = 35). Second, we conducted leave-one-participant-out cross-validated classification (penalty parameter = 0.0001) on the selected participants to validate the mnemonic state classifier and obtained classification accuracy of 60% which is significantly above chance (*t* _34_ = 6.1507, *p <* 0.0001), indicating that the cross-subject mnemonic state classifier is able to distinguish encoding and retrieval states. Finally, we applied the cross-subject mnemonic state classifier to the Phase 2 trials of E1 and E2, and to the test phase trials of E3, specifically in 100 ms intervals from 500 ms pre-response to 500 ms post response. We extracted classifier evidence, the logit-transformed probability that the classifier assigned a given Phase 2 trial a label of encoding or retrieval. This approach provides a trial-level estimate of retrieval state evidence during hit and correct rejection trials.

### Statistical Analyses

We used an independent samples *t*-test to assess the difference in correct rejection (CR) rate between E1 and E2. We used mixed effects ANOVAs and *t*-tests to assess the effect of experiment (E1, E2), response (hit, CR) and time interval on memory state evidence. We used a mixed effects ANOVA and *t*-tests to assess the effect of experiment (E2, E3) and response (hit, CR) on memory state evidence during the pre-response interval. We used a repeated measures ANOVA and *t*-tests to assess the effect of response confidence (high, low) and time interval on memory state evidence for hits in E1. We used false discovery rate (FDR) to correct for multiple comparisons for post-hoc *t*-tests (Benjamini & Hochberg, 1995).

### Code Accessibility

All raw, de-identified data and the associated experimental and analysis codes used in this study will be made available via the Long Term Memory Lab Website upon publication.

## Results

### Top-down goals modulate test phase reaction times

Our general expectation is that the task instructions to either recognize old items (E1) or detect new items (E2) will impact the processes in which participants engage when responding to the test stimuli. Such processing differences may yield accuracy dissociations between experiments. Based on our pilot data, we expected participants in E2 to be more conservative, whereby their CR rates would be lower than those in E1. However, we did not find a significant difference in CR rates (*t* _70_ = 0.3977, *p* = 0.6921, *d* = 0.0951) between E1 (M = 0.8141, SD = 0.1499) and E2 (M = 0.8273, SD = 0.1268; Figure 2A).

**Figure 2.**
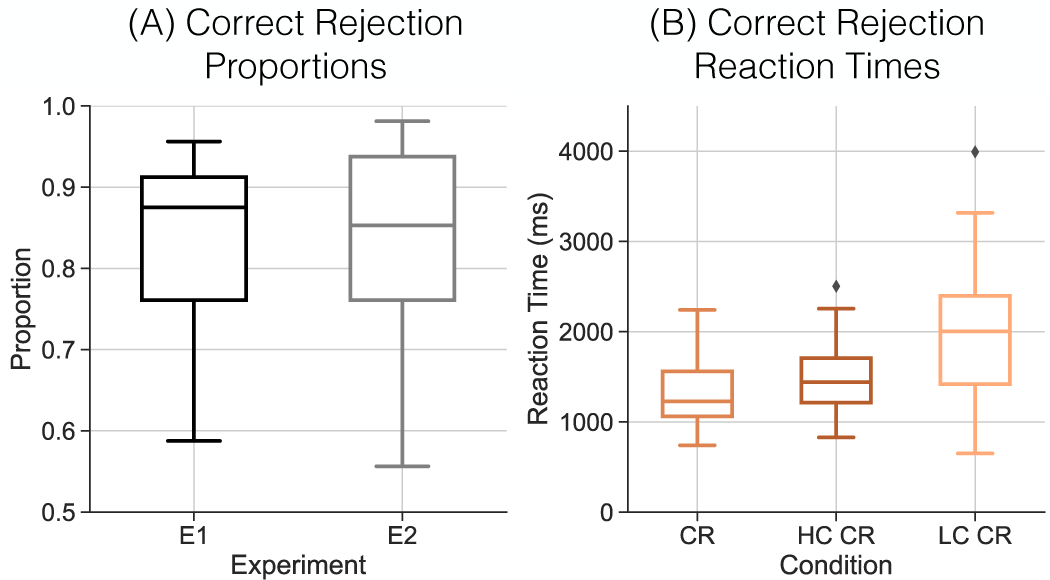
Test Phase Instructions Modulate Correct Rejection Reaction Times. **(A)** Correct rejection (CR) proportions for E1 (black) and E2 (grey). We do not find a significant difference in CR rates across experiments. **(B)** Reaction times (RTs) for correct rejections. E2 CRs are shown in orange. We divided CRs in E1 into high confident (HC, dark orange) and low confident (LC, light orange) responses. We find faster RTs for E2 CRs compared to E1 HC CRs. Box-and-whisker plots show median (center line), upper and lower quartiles (box limits), 1.5x interquartile range (whiskers) and outliers (diamonds).

As the lack of difference in CR rates is difficult to interpret, we conducted an exploratory analysis of reaction times (RTs). Because participants in E1 made confidence judgments whereas participants in E2 did not, we divided response types into three conditions, CRs (E2), high confidence CRs (E1, responses of “1”) and low confidence CRs (E1, responses of “2”; Figure 2B). We find that CRs in E2 (M = 1302 ms, SD = 328.1 ms) are significantly faster than high confidence CRs in E1 (M = 1479 ms, SD = 379.6 ms; *t* _70_ = 2.080, *p* = 0.0411, *d* = 0.4973). These findings demonstrate that the manipulated instruction demands impact recognition memory responses. Specifically, matched top-down instructions (“detect new”) and bottom-up factors (lures), may facilitate responses, hence the faster RTs for CRs in E2 compared to E1.

### Top-down goals modulate retrieval state engagement

Our central goal was to test the hypothesis that top-down and bottom-up factors do not interact and specifically that top-down demands fully engage the retrieval state. We consider stimulus repetition to be a bottom-up factor, here represented by the memory response whereby hits constitute a repetition and CRs do not. We expect hits to induce a retrieval state in both E1 and E2. We consider test phase instructions to be a top-down factor. We expect that the demand to recognize old items (E1) will induce a retrieval state, *regardless of the actual probe*, whereas the demand to detect new items (E2), will not induce a retrieval state. To the extent that these top-down and bottom-up factors do not interact, we expect to find differential levels of retrieval state evidence specifically for CRs between E1 and E2 with no difference in retrieval state evidence for hits. Alternatively, if top-down and bottom-up factors interact, we expect to find greater retrieval state evidence for hits in E1, whereby the combination of explicit demand to retrieve coupled with automatic repetition-driven retrieval will produce greatest engagement of the retrieval state.

To test our hypothesis, we applied a cross-study classifier to the response-locked whole-brain spectral data in each experiment and extracted retrieval state evidence from 500 ms preceding to 500 ms following the response (Figure 3). Following our pre-registration, we conducted a 2×2×10 mixed effects ANOVA with factors of response (hit, CR), experiment (E1, E2) and time interval. We do not find a significant main effect of experiment (*F* _1,70_ = 0.0778, *p* = 0.7812, *η_p_*^2^= 0.0011) or response type (*F* _1,70_ = 0.1511, *p* = 0.6987, *η_p_*^2^= 0.0022). We find a significant main effect of time interval (*F* _9,630_ = 4.644, *p <* 0.0001, *η_p_*^2^= 0.0622), with generally more retrieval state evidence around the time of the response. We do not find a significant interaction between response type and time (*F* _9,630_ = 0.8996, *p* = 0.5251, *η_p_*^2^= 0.0127), response type and experiment (*F* _1,70_ = 1.006, *p* = 0.3193, *η_p_*^2^= 0.3193), or time and experiment (*F* _9,630_ = 1.827, *p* = 0.0604, *η_p_*^2^= 0.0254). We find a significant three-way interaction between response type, time, and experiment (*F* _9,630_ = 1.935, *p* = 0.0446, *η_p_*^2^= 0.0269). We performed post-hoc two sample *t*-tests comparing E1 and E2 at each time point separately for CRs and hits. We find significantly greater retrieval state evidence during E1 compared to E2 CRs in the −500 to −400 ms and −400 to −300 ms intervals (−500 to −400: *t* _70_ = 3.676, *p* = 0.0005, *d* = 0.8788; −400 to −300: *t* _70_ = 2.922, *p* = 0.0047, *d* = 0.6984; FDR corrected). All other post-hoc comparisons did not survive multiple comparisons correction (*t* s < 1.450, *p*s > 0.1514).

**Figure 3.**
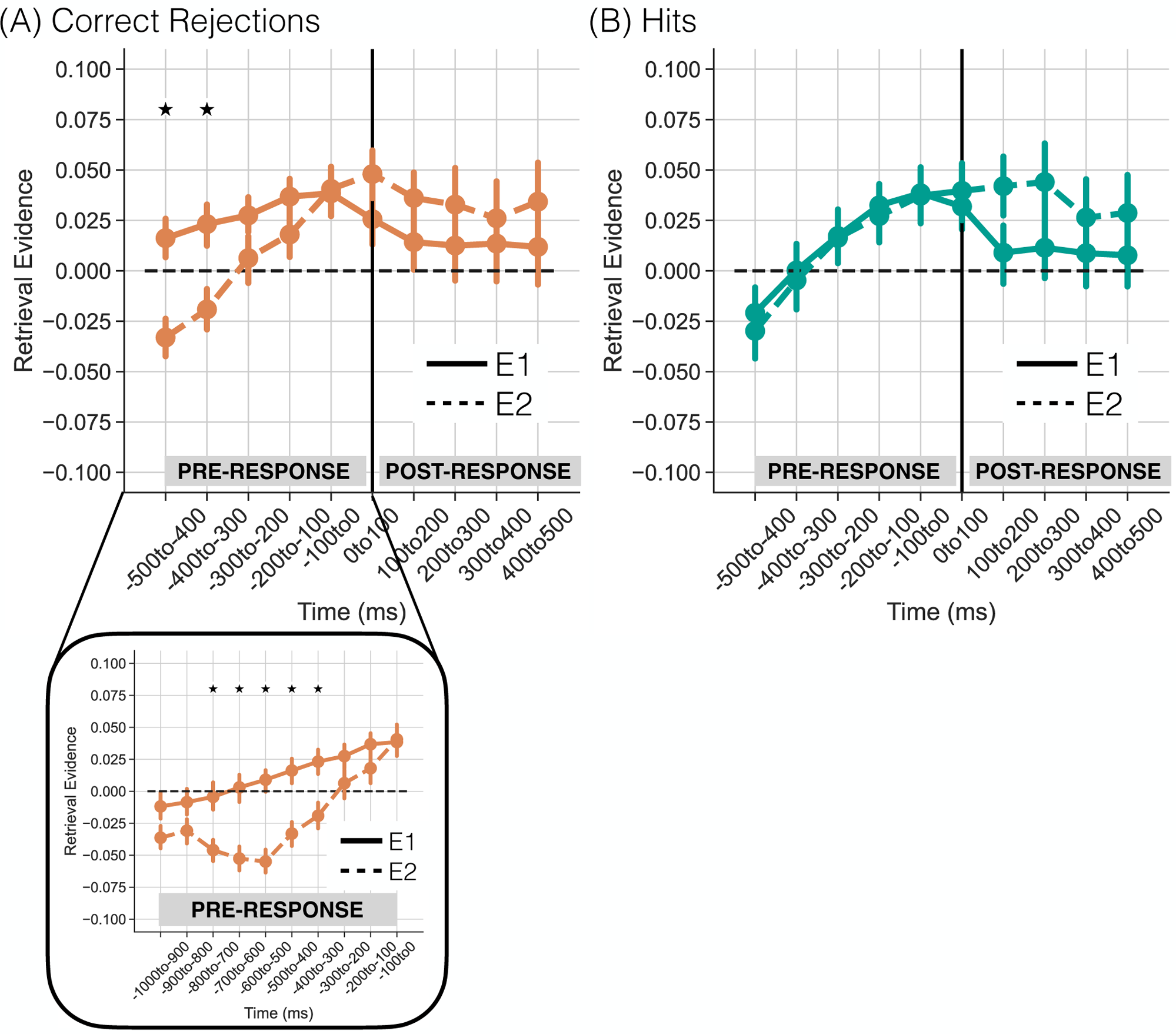
Top-Down Demands Modulate Retrieval State Engagement. We applied a cross-study classifier to response-locked test phase data to measure retrieval state evidence separately for correct rejections (CRs) and hits in E1 (solid) and E2 (dashed). Positive y-axis values indicate greater retrieval state evidence. The vertical line at time 0to100ms indicates the onset of the response. **(A)** Retrieval state evidence preceding the response is significantly greater for CRs in E1 compared to E2. *Lower panel.* When we extend the pre-response interval, we find greater retrieval state evidence for E1 compared to E2 in the 800 to 300 ms preceding the response. **(B)** There is no significant difference in retrieval state evidence for hits across experiments. Stars reflect paired *t*-tests that survived FDR correction. Error bars reflect standard error of the mean.

We had expected that top-down demands could dissociate the two experiments early relative to the response onset and our initial finding of significant dissociations in retrieval state evidence in the 500-300 ms preceding the response is consistent with this expectation. Our next step was to directly test the specificity of the timing of this effect. We averaged retrieval state evidence across the first two time intervals (−500 to −400 and −400 to −300 ms) and across the remaining time intervals (from −300 to 500 ms, relative to response onset). We conducted a 2×2 mixed effects ANOVA with factors of time interval (early, late) and experiment (E1, E2). We do not find a significant main effect of experiment (*F* _1,70_ = 2.058, *p* = 0.156, *η_p_*^2^= 0.0286). We find a significant main effect of time interval (*F* _1,70_ = 6.258, *p* = 0.0147, *η_p_*^2^= 0.0821), driven by greater retrieval state evidence in the late (M = 0.0264, SD = 0.0606) compared to early (M = −0.0032, SD = 0.0605) time interval. We find a significant interaction between time interval and experiment (*F* _1,70_ = 6.113, *p* = 0.0159, *η_p_*^2^= 0.0803), driven by a greater dissociation between experiments in the early compared to late interval. This robust dissociation in time between experiments is consistent with the expectation that top-down demands should engage the retrieval state early relative to response onset.

As it is possible that the observed dissociations in CRs extend even earlier, prior to the *a priori* selected pre-response window of −500 to 0 ms, we conducted an exploratory analysis in which we extended the pre-response interval and measured retrieval state evidence during E1 and E2 CRs for the 1000 ms preceding the response (Figure 3A, lower panel). We conducted a 2×10 mixed effects ANOVA with factors of experiment (E1, E2) and time interval. We find a significant main effect of experiment (*F* _1,70_ = 11.66, *p* = 0.0011, *η_p_*^2^= 0.1428), driven by greater retrieval state evidence for E1 compared to E2. We find a significant main effect of time interval (*F* _9,630_ = 20.37, *p <* 0.0001, *η_p_*^2^= 0.2254), driven by greater retrieval state evidence immediately preceding the response. We find a significant interaction between experiment and time interval (*F* _9,630_ = 3.534, *p* = 0.0003, *η_p_*^2^= 0.0481), driven by greater retrieval state evidence for E1 compared to E2 for the intervals spanning 800 to 300 ms preceding the response (Table 1; FDR corrected). These results show that the retrieval state is engaged several hundred milliseconds prior to a CR specifically when participants’ goal is to recognize old items.

**Table 1.** Post hoc *t*-tests comparing pre-response retrieval state evidence during CRs in E1 and E2.

| Time interval (ms) | E1 vs. E2 |  |  |
| --- | --- | --- | --- |
| | $t_{70}$ | $p$ | $d$ |
| -1000 to -900 | 1.826 | 0.0722 | 0.4364 |
| -900 to -800 | 1.562 | 0.1228 | 0.3733 |
| **-800 to -700** | **2.951** | **0.0043** | **0.7055** |
| **-700 to -600** | **3.904** | **0.0002** | **0.9332** |
| **-600 to -500** | **5.059** | **< 0.0001** | **1.209** |
| **-500 to -400** | **3.676** | **0.0005** | **0.8788** |
| **-400 to -300** | **2.922** | **0.0047** | **0.6984** |
| -300 to -200 | 1.450 | 0.1514 | 0.3467 |
| -200 to -100 | 1.311 | 0.1941 | 0.3134 |
| -100 to 0 | -0.1336 | 0.8941 | 0.0319 |

*Note:* bold values indicate tests that survive FDR correction.

Considering the results for both hits and CRs together, we show that top-down task demands can lead to the recruitment of the retrieval state independent from bottom-up stimulus repetition. That we do not find an additive or over-additive interaction between experiment and response suggests that the combination of the demand to retrieve and stimulus repetition does not further enhance retrieval state engagement beyond either factor alone. Instead, these results suggest that the retrieval state may be engaged any time participants are explicitly directed to retrieve a past experience, regardless of the actual status of the test probe.

### Top-down demands induce the retrieval state in the absence of confidence judgments

Given our motivation to test the hypothesis that a retrieval state engaged by top-down vs. bottom-up factors should differentially relate to the content that is or is not retrieved, participants were instructed to provide a confidence judgment specifically for E1. This manipulation further differentiates E1 from E2 and may thus underlie the observed retrieval state dissociations reported above. Namely, greater retrieval evidence for CRs in E1 may reflect the demand to make a confidence judgment, rather than the demand to recognize old items.

To rule out confidence as an alternative explanation for our findings, we have re-analyzed an independently collected dataset from our lab (Smith et al., 2024, “E3”). In E3, participants performed a traditional old/new recognition memory task on lists of words. The instructions in E3 were identical to those in E1 phases 1 and 2 (Figure 1) except that, critically, there was no demand to provide a confidence judgment. Thus the response options (“old,” “new”) were matched across E2 and E3 and the only difference was the task goal to either detect new items (E2) or recognize old items (E3). We applied our cross study-classifier to the response-locked test phase data in E3.

To the extent that the top-down demand to recognize old items induces a retrieval state, we should again observe an experiment by response interaction, whereby pre-response retrieval evidence is greater for CRs in E3 relative to E2. We averaged retrieval state evidence across the 500 ms pre-response interval (Figure 4). We conducted a 2×2 mixed effects ANOVA with factors of response type (hit, CR), and experiment (E2, E3). We do not find a significant main effect of experiment (*F* _1,72_ = 0.647, *p* = 0.424, *η_p_*^2^= 0.0089) or response type (*F* _1,72_ = 1.459, *p* = 0.2311, *η_p_*^2^= 0.0199). We find a significant interaction between response type and experiment (*F* _1,72_ = 5.311, *p* = 0.0241, *η_p_*^2^= 0.0687). This interaction was driven by numerically greater retrieval evidence for CRs in E3 (M = 0.0254, SD = 0.0503) compared to E2 (M = 0.0025, SD = 0.0543; *t* _72_ = 1.858, *p* = 0.0672, *d* = 0.4377). The difference in retrieval evidence between hits in E3 (M = 0.0055, SD = 0.0448) and E2 (M = 0.0092, SD = 0.0723) was not significant (*t* _72_ = −0.2655, *p* = 0.7914, *d* = 0.0622).

**Figure 4.**
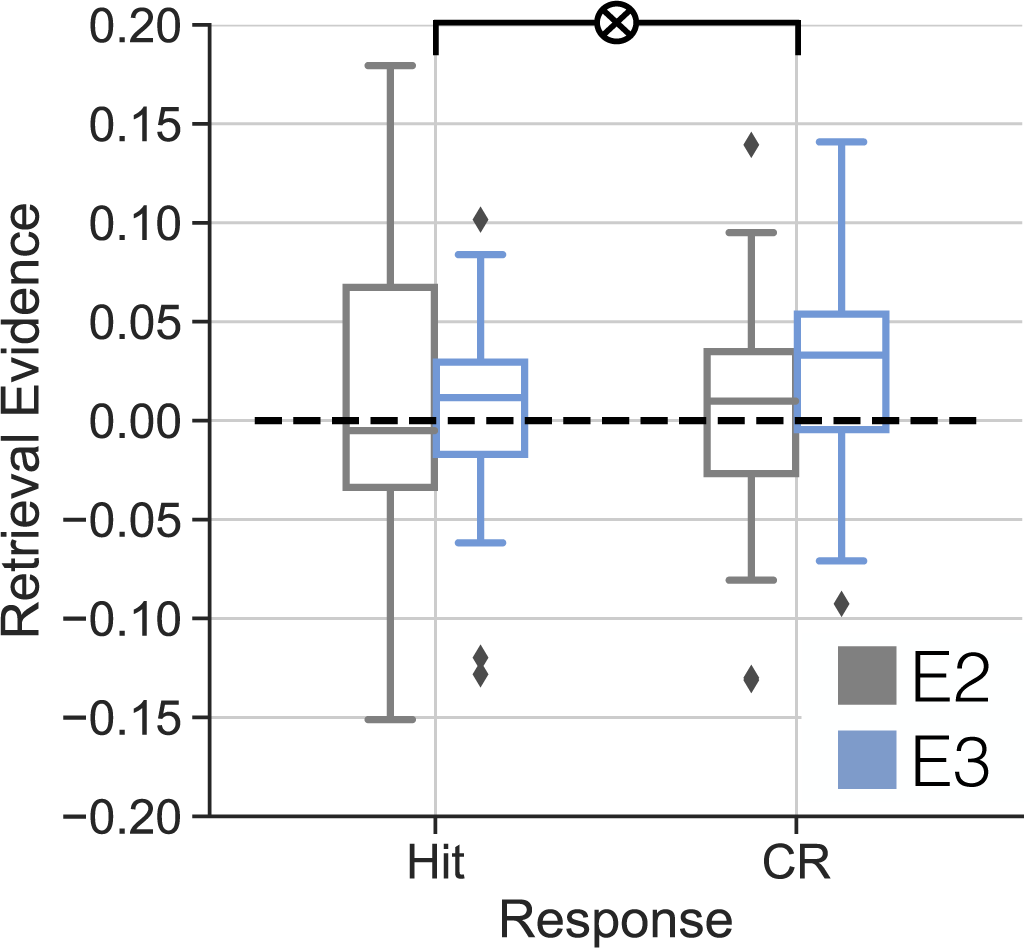
Validation of the Modulation of Top-Down Demands on Retrieval State Engagement. We applied a cross-study classifier to response-locked test phase data in two separate experiments. We extracted average retrieval state evidence in the 500 to 0 ms preceding hits and CRs. In E2 (grey), participants are instructed to detect new items. In E3 (blue), participants are instructed to recognize old items without making confidence judgments. We find a significant response by experiment interaction (*p* = 0.0241). Box-and-whisker plots show median (center line), upper and lower quartiles (box limits), 1.5x interquartile range (whiskers) and outliers (diamonds).

In sum, we validated that the dissociation of retrieval evidence during CRs across experiments is not due to the demand to make a confidence judgment. We find that when participants are instructed to recognize old items – classic recognition memory task instructions – the retrieval state is recruited during CRs, whereas when participants are instructed to detect new items, no such engagement is observed. This result suggests that the top-down goal to retrieve induces a retrieval state even in instances when the probe itself (a lure) may not automatically induce retrieval.

### Top-down retrieval state engagement is modulated by lack of retrieved content

Given that top-down demands engage the retrieval state in the absence of bottom-up factors, our second major goal was to test the extent to which the retrieval state is engaged by the presence or absence of retrieved content. Our initial investigation across experiments E1, E2, and E3 suggests that the retrieval state may be driven by the absence of retrieved content; however, analysis of CRs is ambiguous. Theoretically, there is no retrievable content for CRs, given that the probe is a lure. However, participants may evaluate evidence to make a decision for lures. Hits provide a clearer assessment of retrieved content as the assumption is that participants reinstate study phase content during the test phase in order to make a decision for targets (Danker & Anderson, 2010). The content-present vs. content-absent accounts make opposite predictions for retrieval state engagement during hits made with high vs. low confidence. High confidence responses are thought to be supported by greater reinstatement or overall evidence, relative to low confidence responses (Balsdon, Wyart, & Mamassian, 2020). Therefore, to the extent that the retrieval state tracks retrieved content, there should be more retrieval state evidence during high compared to low confidence hits. Alternatively, to the extent that the retrieval state tracks attention or control required to access a representation, there should be more retrieval state evidence during low compared to high confidence hits. The logic is that less evidence should require more internal attention to support a decision, possibly by prolonging the search or evidence accumulation process (Callaway, Griffiths, Norman, & Zhang, 2023).

We tested the content-present vs. content-absent accounts by assessing retrieval state evidence as a function of confidence specifically for E1 hits (Figure 5). Following our pre-registration, we conducted a 2 × 10 repeated measures ANOVA with factors of response confidence (high, low) and time interval (−500 to 500 in 100 ms intervals, response-locked). We find a significant main effect of response confidence (*F* _1,35_ = 4.556, *p* = 0.0399, *η_p_*^2^= 0.1152) driven by greater retrieval state evidence for low (M = 0.037, SD = 0.0766) compared to high (M = 0.0095, SD = 0.0512) confidence hits. We find a significant main effect of time interval (*F* _9,315_ = 2.404, *p* = 0.012, *η_p_*^2^= 0.0643), driven by greater pre-response compared to post-response retrieval evidence. We find a significant interaction between response confidence and time interval (*F* _9,315_ = 7.597, *p <* 0.0001, *η_p_*^2^= 0.1783), whereby retrieval state evidence is significantly greater pre-response for low (M = 0.0639, SD = 0.0735) compared to high (M = 0.0049, SD = 0.0735) confidence hits (*t* _35_ = 3.697, *p* = 0.0007, *d* = 0.8026). Post-response retrieval state evidence does not significantly differ between low (M = 0.0101, SD = 0.1166) and high (M = 0.0142, SD = 0.072) confidence hits (*t* _35_ = 0.2887, *p* = 0.7745, *d* = 0.0422). We performed post-hoc paired sample *t*-tests comparing high and low confidence hits at each time point. We find significantly greater retrieval state evidence during low compared to high confidence hits in the first three pre-response intervals (−500 to −400: *t* _35_ = 5.079, *p <* 0.0001, *d* = 0.9588; −400 to −300: *t* _35_ = 3.849, *p* = 0.0005, *d* = 0.8432; −300 to −200: *t* _35_ = 3.386, *p* = 0.0018, *d* = 0.7156; FDR corrected). All other post-hoc comparisons did not survive multiple comparisons correction (*t*’s < 1.743, *p*’s > 0.0901). Together, these results provide support for the content-absent account whereby the retrieval state is more strongly engaged during trials in which less evidence is available to make a decision.

**Figure 5.**
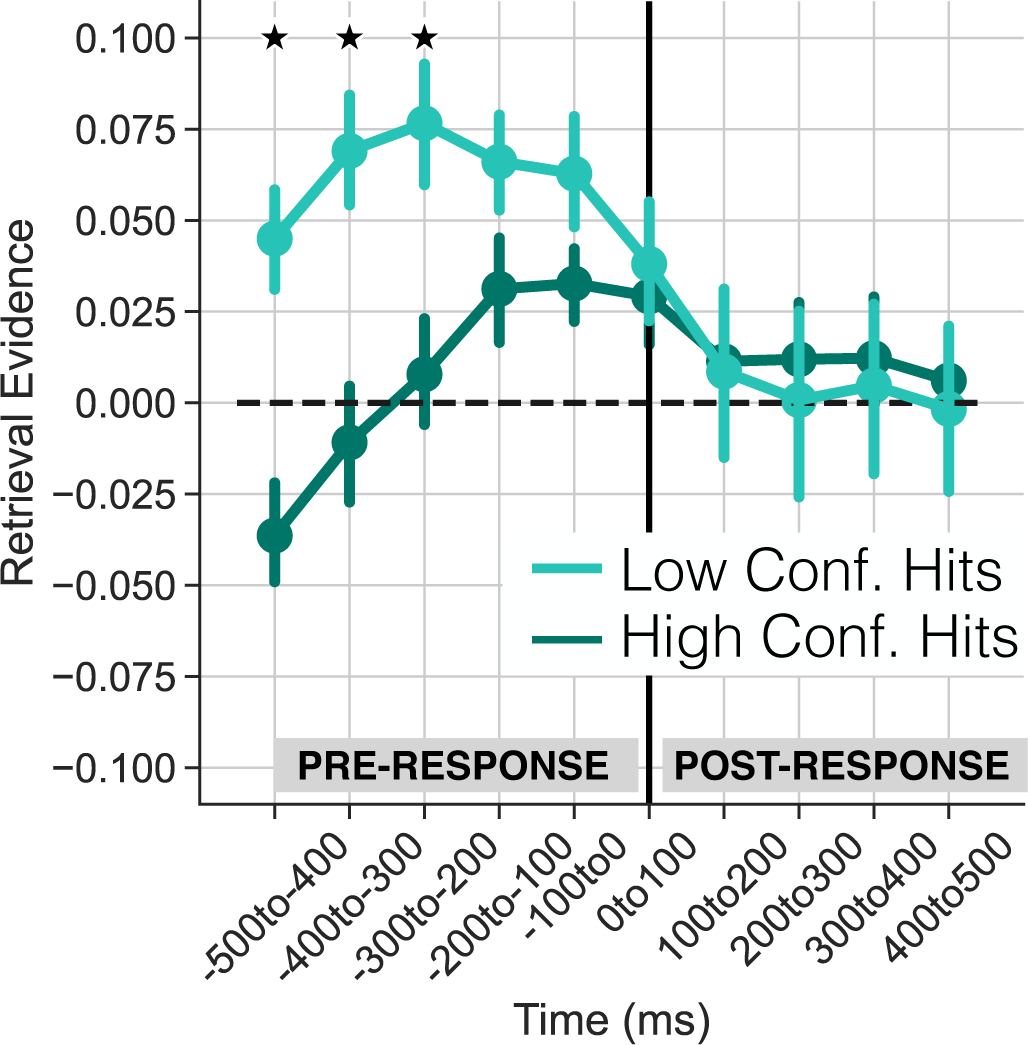
Confidence during Successful Retrieval Modulates Retrieval State Engagement. We applied a cross-study classifier to response-locked test phase data in E1 across ten time intervals from 500 ms preceding to 500 ms following the response. We measured retrieval state evidence separately for high confidence hits (dark teal) and low confidence hits (light teal). We find greater retrieval state evidence for low compared to high confidence hits in the 500 to 200 ms intervals preceding the response (*p*’s < 0.0018). Stars reflect paired *t*-tests that survived FDR correction. Error bars reflect standard error of the mean.

## Discussion

The aim of this study was to identify the joint impact of bottom-up and top-down factors on retrieval state engagement. We conducted two independent recognition memory experiments in which we manipulated top-down goals to either recognize old items (E1) or detect new items (E2). We included a third experiment (E3) to control for possible effects of making a confidence judgment. We recorded scalp EEG and used a cross-study decoding approach (Long, 2023) to measure response-locked retrieval state engagement during the test phase of each experiment. Retrieval state engagement was greater during correct rejections (CRs) when participants’ goal was to recognize old items (E1 and E3) compared to when participants’ goal was to detect new items (E2). Furthermore, retrieval state engagement was greater for low, compared to high confidence hits in E1, when retrieved content is presumably low. Together, these findings suggest that the explicit top-down demand to remember the past recruits the retrieval state independent from bottom-up factors such as stimulus repetition. This intentional retrieval state engagement may reflect internally directed attention deployed to accomplish the task goal of recognizing old items.

We find evidence for a top-down driven retrieval state that is independent from bottom-up factors. Whereas correctly identifying study items as old (hits) recruits the retrieval state regardless of task goals, correct rejection of novel lures (CRs) only recruits the retrieval state when the task goal is to recognize old items. Similarly, we find that retrieval state evidence is greater when participants make low compared to high confidence hits when directed to recognize old items (E1). These findings are consistent with the conceptualization of the retrieval mode as an intentional state that is a precursor to, but separate from, retrieval success (Tulving, 1983). The notion is that one must enter a retrieval state in order to interpret a stimulus as a retrieval cue (Rugg & Wilding, 2000). Thus, in the present study, because participants were explicitly instructed to recognize old items in E1, they engaged a retrieval state and treated every test probe as a retrieval cue. That retrieval state engagement is not driven by retrieval success is supported by the E1 confidence contrast. If the retrieval state specifically reflected retrieval success, we would not expect to find a confidence-based modulation, as both trials constitute retrieval success. If anything, we might expect greater retrieval state evidence for high compared to low confidence hits, whereby stronger representations produce greater retrieval state engagement. Instead, we find greater retrieval state evidence for low compared to high confidence hits which may reflect differences in memory search or evidence accumulation across the two conditions (Balsdon et al., 2020; Callaway et al., 2023; Lee, Daunizeau, & Pezzulo, 2023). Together, these results are consistent with the interpretation that the retrieval state is engaged via top-down demands in service of the attempt to access an internally stored representation.

Our results suggest that the retrieval state is differentially engaged based on task goals. Unlike prior retrieval state work in which episodic retrieval is contrasted with non-episodic tasks such as semantic processing (Morcom & Rugg, 2002; Williams & Wilding, 2019) and perceptual processing (Evans, Williams, & Wilding, 2015), in the present study we used two very closely matched test phase goals. According to existing behavioral findings (Brainerd, Bialer, Chang, & Upadhyay, 2021), “recognize old” and “detect new” are not simply inverses of one another. Instead, the specific goal that participants are given informs how they interact with test probes and will lead to differential processing of targets vs. lures depending on whether they are tasked with recognizing old items vs. detecting new items. Such task-based dissociations in stimulus processing are consistent with neural evidence showing that shifts in top-down goals can change how stimuli are represented in the brain (Aly & Turk-Browne, 2016; Lim, Lang, & Diana, 2023; Long & Kuhl, 2021). Our results extend these findings by showing that not only do task goals impact stimulus representations, but also modulate the processes or states that are engaged in the service of those task goals.

If the retrieval state is only engaged for CRs when participants recognize old items, what processes are engaged when participants are instructed to detect new items in E2? By one account, participants may engage in some form of novelty detection and greater univariate activity for lures vs. targets may reflect a novelty signal (Tulving, Markowitsch, Craik, Habib, & Houle, 1996; Knight, 1996; Grunwald, Lehnertz, Heinze, Helmstaedter, & Elger, 1998; Ranganath & Rainer, 2003; Düzel, Penny, & Burgess, 2010). Potentially, the detection of novelty may induce an encoding state (Patil & Duncan, 2018), governed by acetylcholine release. The design of our independent mnemonic state classifier is such that “negative” retrieval state evidence is equivalent to evidence for an encoding state. We find evidence for encoding state engagement during CRs in E2 in the early time intervals preceding the response, which is consistent with a novelty/encoding interpretation. However, it is important to note that despite the differences leading into CRs, we do find evidence of retrieval state engagement immediately preceding all responses in both E1 and E2. We speculate that this retrieval state response may reflect the decision process, rather than a memory process per se. In prior work, we have found that retrieval state engagement “ramps up” leading to a behavioral decision in a spatial attention task (Long, 2023), suggesting that the retrieval state may be engaged to evaluate a decision in addition to supporting episodic memory retrieval attempts. Future work will be needed to specifically address how memory vs. decision-making processes engage memory brain states.

The top-down induced retrieval state may reflect internal attention engaged in the service of the attempt to retrieve. Both theoretical proposals and computational models suggest that long term memory retrieval reflects a form of internally directed attention (Chun et al., 2011; Long, Kuhl, & Chun, 2018; Logan, Cox, Annis, & Lindsey, 2021). Our interpretation is that the instruction to recognize old items induces participants to turn their mind’s eye inward when evaluating test probes. When more evidence is required to evaluate a probe, more internal attention should be recruited, which is consistent with our finding of greater retrieval state engagement during low compared to high confidence hits. We anticipate that under any circumstances in which participants are given and follow traditional memory task instructions to recognize or remember old items, the retrieval state will be recruited, regardless of the status of the test probe. There may be important individual differences in the ability to engage the retrieval state in the service of task goals, which will in turn impact downstream processing and behavior.

Despite clear evidence that top-down demands engage the retrieval state, we still expect that bottom-up factors play a role in memory state engagement. We observe retrieval state engagement for hits in both E1 and E2, as would be expected from a bottom-up account. Behavioral evidence suggests that old vs. new stimuli may lead the hippocampus into a pattern separating vs. pattern completing state (Duncan et al., 2012; Patil & Duncan, 2018). When two categorically related experiences occur near together in time – compared to far apart in time – there is greater engagement of the retrieval state (Smith et al., 2022). From a purely top-down perspective, far trials should be more difficult to retrieve than near trials, similar to low vs. high confidence hits in E1, meaning that there should be more retrieval evidence for far than near trials. However, the opposite finding of greater retrieval evidence for near than far trials suggests a bottom-up influence of stimulus features, in this case temporal overlap, on retrieval state engagement. Our interpretation of these prior findings in light of the current findings is that the retrieval state can be induced automatically via bottom-up inputs, but that these effects are likely to be largely masked by top-down factors and will only emerge either in the absence of top-down demands or when bottom-up factors are highly salient. Our manipulation of bottom-up factors in the present study was relatively modest, with targets that were only repeated once, having been presented during study and during test. Thus, with more substantial manipulations – more repetitions, higher similarity to other stimuli, etc. – we might find greater influences of bottom-up factors on the retrieval state. An exciting future direction will be to investigate the full range of bottom-up factors that can induce retrieval state engagement, especially in the absence of top-down factors.

Taken together, our findings indicate that the retrieval state is engaged in the service of the top-down goal to recognize old items. The retrieval state may thus be recruited in response to any demand to direct attention internally, regardless of whether stimuli themselves have previously been encountered. These results advance our understanding of the interplay between top-down and bottom-up factors on memory state engagement.

## Acknowledgments

This work was supported by a grant from the National Institutes of Health (NINDS R01 NS132872, PI: NML).

## References

Aly, M., & Turk-Browne, N. B. (2016). Attention promotes episodic encoding by stabilizing hippocampal representations. Proceedings of the National Academy of Sciences of the United States of America, 113(4), E420–E429.

Balsdon, T., Wyart, V., & Mamassian, P. (2020). Confidence controls perceptual evidence accumulation. Nature Communications, 11(1753).

Benjamini, Y., & Hochberg, Y. (1995). Controlling the false discovery rate: a practical and powerful approach to multiple testing. Journal of the Royal Statistical Society. Series B (Methodological), 57 (1), 289–300.

Brainerd, C. J., Bialer, D. M., Chang, M., & Upadhyay, P. (2021). A fundamental asymmetry in human memory: Old = not-new and new = not-old. Journal Experimental Psychology: Learning, Memory, and Cognition.

Cabeza, R., Mazuz, Y. S., Stokes, J., Kragel, J. E., Woldorff, M. G., Ciaramelli, E.,… Moscovitch, M. (2011). Overlapping parietal activity in memory and perception: evidence for the attention to memory model. Journal of Cognitive Neuroscience, 23(11), 3209–3217.

Callaway, F., Griffiths, T. L., Norman, K. A., & Zhang, Q. (2023). Optimal metacognitive control of memory recall. Psychological Review.

Castellanos, N. P., & Makarov, V. A. (2006). Recovering EEG brain signals: Artifact suppression with wavelet enhanced independent component analysis. Journal of Neuroscience Methods, 158(2), 300–312.

Chun, M. M., Golomb, J. D., & Turk-Browne, N. B. (2011). A taxonomy of external and internal attention. Annual Review of Psychology, 62, 73–101.

Danker, J. F., & Anderson, J. R. (2010). The ghosts of brain states past: Remembering reactivates the brain regions engaged during encoding. Psychological Bulletin, 136(1), 87–102.

Duncan, K. D., Sadanand, A., & Davachi, L. (2012). Memory’s penumbra: Episodic memory decisions induce lingering mnemonic biases. Science, 337 (6093), 485–487.

Düzel, E., Penny, W. D., & Burgess, N. (2010). Brain oscillations and memory. Current Opinion in Neurobiology, 20(2), 143–149.

Evans, L. H., Williams, A. N., & Wilding, E. L. (2015). Electrophysiological evidence for retrieval mode immediately after a task switch. NeuroImage, 108, 435–440.

Friendly, M., Franklin, P. E., Hoffman, D., & Rubin, D. C. (1982). The Toronto Word Pool: Norms for imagery, concreteness, orthographic variables, and grammatical usage for 1,080 words. Behavior Research Methods & Instrumentation, 14, 375–399.

Grunwald, T., Lehnertz, K., Heinze, H. J., Helmstaedter, C., & Elger, C. E. (1998). Verbal novelty detection within the human hippocampus proper. Proceedings of the National Academy of Sciences of the United States of America, 95, 3193–3197.

Harris, K. D., & Thiele, A. (2011). Cortical state and attention. Nature Reviews Neuroscience, 12(9), 509–523.

Herron, J. E., & Wilding, E. L. (2004). An electrophysiological dissociation of retrieval mode and retrieval orientation. NeuroImage, 22, 1554–1562.

Herron, J. E., & Wilding, E. L. (2006). Brain and behavioral indices of retrieval mode. NeuroImage, 32, 863–870.

Hong, Y., Smith, D. E., Moore, I. L., & Long, N. M. (2023). Spatiotemporal dynamics of memory encoding and memory retrieval states. Journal of Cognitive Neuroscience, 35(9).

Knight, R. T. (1996). Contribution of human hippocampal region to novelty detection. Nature, 383, 256–259.

Kuhl, B. A., & Chun, M. M. (2014). Successful remembering elicits event-specific activity patterns in lateral parietal cortex. The Journal Of Neuroscience, 34(23), 8051–8060.

Lee, D. G., Daunizeau, J., & Pezzulo, G. (2023). Evidence or confidence: What is really monitored during a decision? Psychonomic Bulletin & Review.

Lepage, M., Ghaffar, O., Nyberg, L., & Tulving, E. (2000). Prefrontal cortex and episodic memory retrieval mode. Proceedings of the National Academy of Sciences of the United States of America, 97 (1), 506–511.

Lim, Y.-L., Lang, D. J., & Diana, R. A. (2023). Cognitive tasks affect the relationship between representational pattern similarity and subsequent item memory in the hippocampus. NeuroImage, 227.

Logan, G. D., Cox, G. E., Annis, J., & Lindsey, D. R. B. (2021). The episodic flanker effect: Memory retrieval as attention turned inward. Psychological Review, 128(3), 397–445.

Long, N. M. (2023). The intersection of the retrieval state and internal attention. Nature Communications, 14(3861). doi: 10.1038/s41467-023-39609-9

Long, N. M., & Kahana, M. J. (2015). Successful memory formation is driven by contextual encoding in the core memory network. NeuroImage, 119, 332–337.

Long, N. M., & Kuhl, B. A. (2018). Bottom-up and top-down factors differentially influence stimulus representations across large-scale attentional networks. Journal of Neuroscience, 38(10), 2495–2504.

Long, N. M., & Kuhl, B. A. (2019). Decoding the tradeoff between encoding and retrieval to predict memory for overlapping events. NeuroImage, 201.

Long, N. M., & Kuhl, B. A. (2021). Cortical representations of visual stimuli shift locations with changes in memory states. Current Biology, 31(5).

Long, N. M., Kuhl, B. A., & Chun, M. M. (2018). Memory and attention. In J. T. Wixted (Ed.), Steven’s handbook of experimental psycholgoy and cognitive neuroscience (Fourth ed.). John Wiley & Sons, Inc.

Morcom, A. M., & Rugg, M. D. (2002). Getting ready to remember: the neural correlates of task set during recognition memory. NeuroReport, 13(1).

Nolan, H., Whelan, R., & Reilly, R. (2010). Faster: fully automated statistical thresholding for eeg artifact rejection. Journal of neuroscience methods, 192(1), 152–162.

O’Reilly, R. C., & McClelland, J. L. (1994). Hippocampal conjunctive encoding, storage, and recall: avoiding a trade-off. Hippocampus, 4(6), 661–682.

Patil, A., & Duncan, K. (2018). Lingering cognitive states shape fundamental mnemonic abilities. Psychological Science, 29(1), 45–55.

Ranganath, C., & Rainer, G. (2003). Neural mechanisms for detecting and remembering novel events. Nature Reviews Neuroscience, 4(3), 193–202.

Rugg, M. D., & Wilding, E. L. (2000). Retrieval processing and episodic memory. Trends in Cognitive Sciences, 4(3), 108–115.

Smith, D. E., Moore, I. L., & Long, N. M. (2022). Temporal context modulates encoding and retrieval of overlapping events. Journal of Neuroscience, 42(14), 3000–3010.

Smith, D. E., Wheelock, J. R., & Long, N. M. (2024). Response-locked theta dissociations reveal potential feedback signal following successful retrieval. bioRxiv 575166. doi: 10.1101/2024.01.11.575166

Tulving, E. (1983). Elements of episodic memory. New York: Oxford.

Tulving, E., Markowitsch, H. J., Craik, F. I., Habib, R., & Houle, S. (1996). Novelty and familiarity activations in pet studies of memory encoding and retrieval. Cerebral Cortex, 6(1), 71–79.

Williams, A. N., & Wilding, E. L. (2019). On the sensitivity of event-related potentials to retrieval mode. Brain and Cognition, 135.

Yassa, M. A., & Stark, C. E. L. (2011). Pattern separation in the hippocampus. Trends in Neurosciences, 34(10), 515–525.

